# H3K9me3 as a gatekeeper of lineage-specific enhancers in embryonic progenitor cells

**DOI:** 10.64898/2026.05.03.722466

**Authors:** Kenji Ito, Greg Donahue, Takeshi Katsuda, Kenji Kamimoto, Kenneth S. Zaret

**Affiliations:** Institute for Regenerative Medicine, Department of Cell and Developmental Biology, Penn Epigenetics Institute Perelman School of Medicine, University of Pennsylvania, Philadelphia, PA, USA; Department of Chemical System Engineering, Graduate School of Engineering, The University of Tokyo; Department of Systems Biomedical Science, Research Institute for Microbial Diseases, The University of Osaka, Suita, Osaka, Japan

**Keywords:** H3K9me3, heterochromatin, repression, enhancers, gene expression

## Abstract

While many studies of developmental control have focused on gene activation, less is known about the extent to which regulatory programs are actively repressed in progenitor cells. We previously showed that trimethylation of histone H3 at lysine 9 (H3K9me3) is a repressive mark that is remodeled on protein-coding genes when endodermal progenitors transition to liver and pancreatic β cell fates. Yet whether H3K9me3 is dynamic at promoters and enhancers has not been determined. Here we find that promoters of liver-specific genes are strongly enriched for H3K9me3 in undifferentiated progenitors, whereas such enrichment is not observed at promoters of more broadly expressed liver genes. We further show that enhancers specific to differentiated tissues—including liver, islet, and cerebral cortex—are strongly enriched for H3K9me3 in their corresponding tissue stem and progenitor cells. In hepatoblasts, H3K9me3 contributes to maintaining the undifferentiated state by restricting FOXA2 and HNF4α from binding to most enhancers, while there remain thousands of H3K9me3-marked enhancers where the factors are not restricted from binding. Our findings illustrate how H3K9me3-mediated heterochromatinization can restrict transcription factor engagement in progenitor cells to prevent inappropriate activation during early development. H3K9me3 at enhancers that allow transcription factor binding may reflect developmental competence.

## Introduction

Somatic cell identity is progressively established during development through the integration of extrinsic signaling cues and the coordinated activation of cell type–specific transcriptional programs (Zaret 2008; Hirabayashi and Gotoh 2010). Once established, cellular identity is maintained by the interplay between transcription factor networks and epigenetic regulation of the genome, particularly at distal regulatory elements such as enhancers (Bulger and Groudine 2011; Spitz and Furlong 2012; Whyte et al. 2013; Hnisz et al. 2016). Trimethylation of histone H3 at lysine 9 (H3K9me3) has been implicated in safeguarding cell identity by repressing lineage-inappropriate gene expression (Soufi et al. 2012; Becker et al. 2016; Becker et al. 2017; Nicetto and Zaret 2019; Methot et al. 2021; Padeken et al. 2022). Previous studies have shown that hepatocyte and pancreatic beta-cell differentiation is accompanied by a dynamic loss of H3K9me3 at mouse hepatocyte-specific gene loci, coinciding with transcriptional activation (Nicetto et al. 2019). However, that study, like most others, focused on H3K9me3 across the transcribed region of genes. How H3K9me3 may be differentially distributed at promoters and enhancers over developmental time and the functional significance of such redistribution for cell identity transitions has not been evaluated.

To address this gap, we performed integrative bioinformatic analyses across multiple developmental systems. First, we identified tissue-specific cis-regulatory elements and examined H3K9me3 dynamics at these regions across developmental stages. To identify candidate factors that contribute to dynamic H3K9me3 changes, we used mass spectrometry data of HDAC3-associated proteins generated in differentiated hepatocytes and assessed their expression levels in undifferentiated cells using published RNA-seq data. Finally, we integrated ChIP-seq data generated across multiple stages of liver development and found that H3K9me3 contributes to maintaining the undifferentiated state through distinct molecular mechanisms, depending on enhancer class.

## Results

### Adult liver–specific promoters are selectively marked by H3K9me3 in undifferentiated progenitors

To examine how H3K9me3 contributes to lineage control during liver development, we analyzed H3K9me3 ChIP-seq datasets spanning differentiation from definitive endoderm to mature hepatocytes (Fig. 1A). Our previous work identified dynamic H3K9me3 remodeling at hepatocyte gene loci during this process (Nicetto et al. 2019). While promoter chromatin states are generally stable across cell types, enhancer chromatin states are highly tissue-restricted and developmentally regulated (Thurman et al. 2012; Roadmap Epigenomics et al. 2015), suggesting that enhancer repression may represent a critical regulatory layer during lineage specification.

**Fig. 1.**
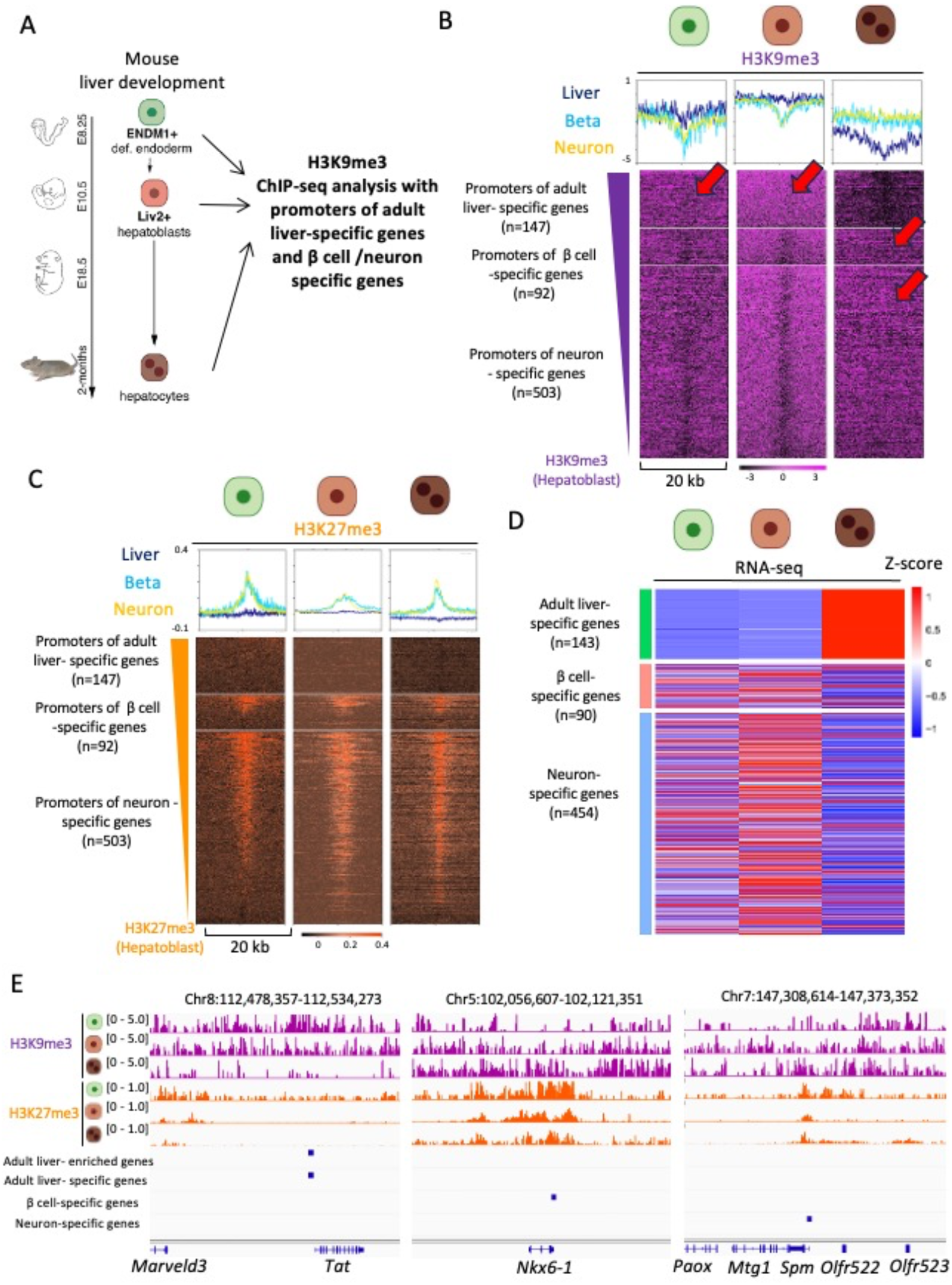
Adult liver-specific promoters are highly marked by H3K9me3 in undifferentiated endoderm and hepatoblasts. A. Schematic illustration of the developmental stages and cell types analyzed in this figure. B. H3K9me3 heatmaps centered on promoters of adult liver-specific genes (top), β-cell-specific genes (middle), and neuron-specific genes (bottom) across liver developmental stages. C. H3K27me3 heatmaps centered on promoters of adult liver-specific genes (top), β-cell-specific genes (middle), and neuron-specific genes (bottom) across liver developmental stages. D. Heatmap showing expression of liver-specific genes (top), β-cell-specific genes (middle) and neuron-specific genes (bottom) during liver development based on published RNA-seq data (Nicetto et al., 2019). E. Genome browser view of the adult liver-specific *Tat* locus (left), the β-cell-specific *Nkx6-1* locus (center), and the neuron-specific *Spm* locus (right) showing H3K9me3 and H3K27me3 dynamics during liver differentiation from definitive endoderm to hepatocytes. For heatmap visualization, averaged H3K9me3 and H3K27me3 ChIP–seq signal from biological replicates at each developmental stage were used. H3K9me3 and H3K27me3 signals represent Input-subtracted tracks. The same processed signal tracks used in Fig.1B and 1C were used for IGV visualization in Fig. 1E. H3K9me3 and H3K27me3 heatmaps in Fig.1B and 1C are sorted based on signal intensity in hepatoblasts.

In a previous study, we identified genes that are highly expressed in differentiated hepatocytes, but not in endoderm and hepatoblasts, during mouse liver development from endoderm by performing bulk RNA-seq (Cluster #13 in Nicetto et al., 2019; Table S1; hereafter referred to as adult liver-enriched genes). We first examined the H3K9me3 state around the promoters of adult liver-enriched genes (Table S2) and found an absence of H3K9me3 throughout liver development from endoderm, even though these genes are silenced in endoderm and hepatoblasts (Fig. S1A and S1B). Adult liver-specific genes, which are exclusively expressed in hepatocytes but not in other adult tissues, were further identified from promoters of the adult liver-enriched genes that overlap with adult liver-specific RNA Pol II ChIP-seq peaks (Fig. S1C and Table S3 and S4; see also Experimental Procedures). GO analysis of adult liver-specific genes showed that genes related to metabolic functions are significantly enriched (Fig. S1D).

As controls, promoters of genes specific to differentiated β cells and neurons, which were identified from previously published datasets (Benner et al. 2014; Sanosaka et al. 2017) (Table S5 and S6), were used. Comparison of H3K9me3 states around promoters of liver-specific genes throughout liver development showed that these promoters are marked by H3K9me3 in endoderm and hepatoblasts, whereas H3K9me3 levels were markedly reduced in differentiated hepatocytes (Fig. 1B, top, Fig. 1E left). Interestingly, H3K9me3 around promoters of the alternative lineages showed inverse dynamics: H3K9me3 levels are reduced in endoderm and hepatoblasts, but these regions are marked by H3K9me3 in differentiated hepatocytes (Fig. 1B, middle and bottom, Fig. 1E center and right). Notably, promoters of differentiated β cell- and neuron-specific genes are highly marked by H3K27me3 throughout liver development (Fig. 1C, middle and bottom, Fig. 1E center and right). In differentiated hepatocytes, promoters of these control lineages are marked by both H3K9me3 and H3K27me3 (Fig. 1B, 1C, and 1E), indicating that the cellular identity of differentiated hepatocytes is reinforced by repressing promoters of alternative lineages through both modifications. Genes specific to β cells and neurons are repressed in differentiated hepatocytes, but their expression is weakly observed in both endoderm and hepatoblasts (Fig. 1D). In contrast, adult liver-specific genes are repressed in endoderm and hepatoblasts without H3K27me3, indicating that the H3K9me3 mark around adult liver-specific genes is more important for their repression than H3K27me3 in the undifferentiated state (Fig. 1D).

Together, these results indicate that the H3K9me3 mark at promoters of lineage-specific genes is predominantly associated with stable gene repression, rather than H3K27me3, and that the role of H3K9me3 in promoter repression of alternative lineages varies depending on the stage of liver development. These findings prompted us to ask whether similar H3K9me3-mediated repression extends to lineage-specific enhancers, which play a central role in cell fate determination.

### Adult liver–specific enhancers are selectively marked by H3K9me3 in undifferentiated progenitors

To extend these findings, we next focused on adult liver-specific enhancer elements.

Adult liver-specific enhancer elements were identified by taking the subset of adult liver specific enhancers (Shen et al. 2012) that overlap with adult liver DNase I hypersensitive sites (Fig. 2A and Table. S7, see also Experimental Procedures). As controls, we compared the data with that from previously reported adult-islet specific enhancers (Tennant et al. 2013) and newly established adult-cortex specific enhancers (Fig. S2A and Table S7, see also Experimental Procedures). Analysis of adult liver-specific enhancers revealed a more pronounced H3K9me3 dynamics: adult liver-specific enhancers were strongly enriched for H3K9me3 in undifferentiated definitive endoderm and hepatoblasts, while H3K9me3 was largely depleted from these enhancers in mature hepatocytes (Fig. 2B). By contrast, adult islet- and cortex-specific enhancers remained highly enriched for H3K9me3 throughout liver development (Fig. 2C), consistent with stable repression of lineage-inappropriate regulatory elements.

**Fig. 2.**
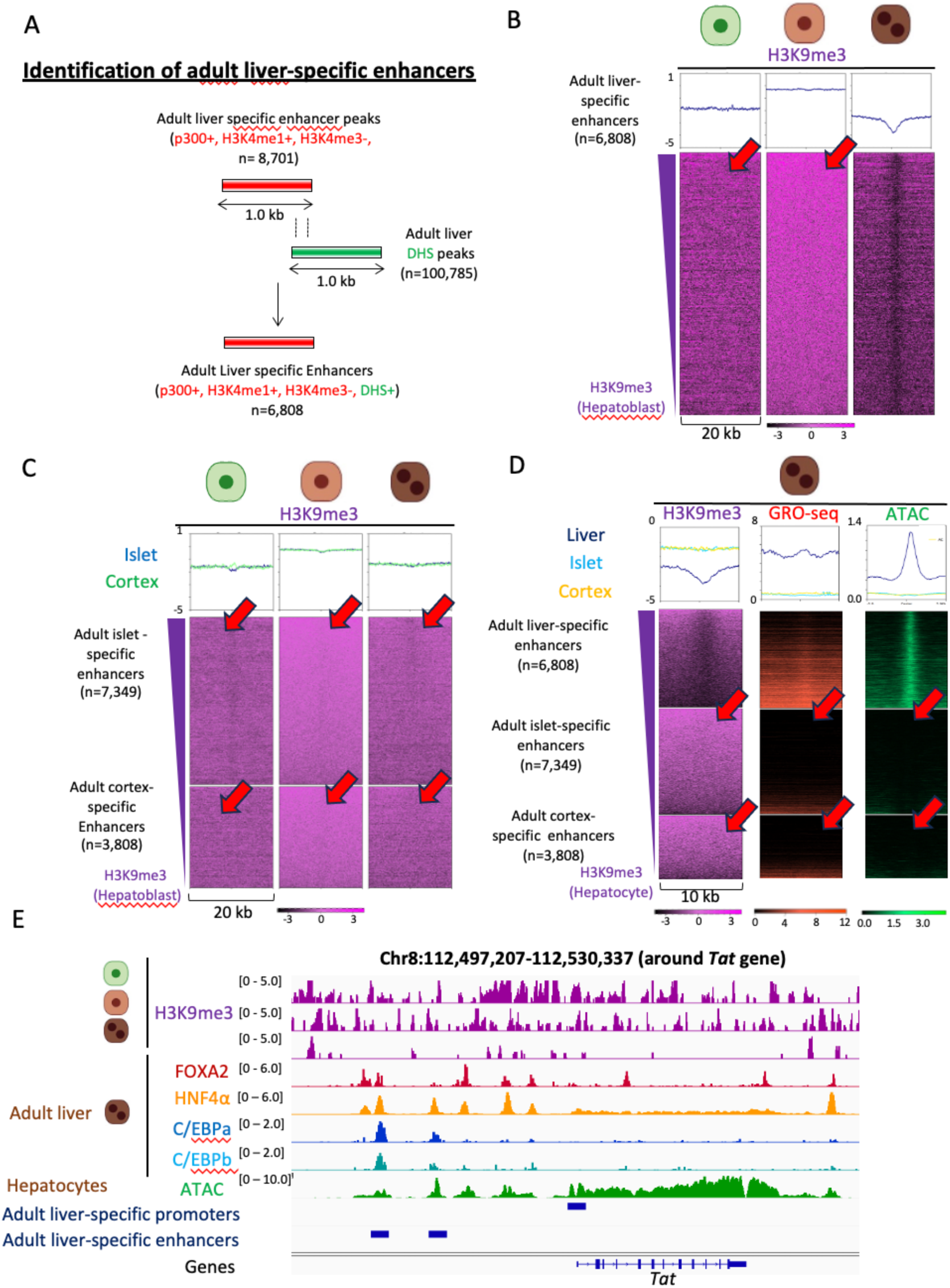
Adult liver-specific enhancers are highly marked by H3K9me3 in undifferentiated endoderm and hepatoblasts. A. Schematic illustration of the identification process for adult liver-specific enhancers. B. H3K9me3 heatmaps centered on adult liver-specific enhancers across liver developmental stages. C. H3K9me3 heatmaps centered on adult islet-specific enhancers (top) and adult cortex-specific enhancers (bottom) across liver developmental stages. D. H3K9me3, GRO-seq, and ATAC-seq heatmaps in adult liver centered on adult liver–specific enhancers (top), adult islet-specific enhancers (middle), and adult cortex-specific enhancers (bottom). Positive- and negative-strand GRO-seq signals were merged prior to analysis. Islet- and cortex-specific enhancers, which retain H3K9me3 in liver, exhibit minimal enhancer-associated nascent transcription and chromatin accessibility compared with adult liver–specific enhancers. E. Genome browser view of the hepatocyte-specific *Tat* locus showing H3K9me3 dynamics during liver differentiation alongside binding of hepatocyte lineage-determining transcription factors and chromatin accessibility (ATAC-seq) in differentiated liver. For heatmap visualization, averaged H3K9me3 ChIP–seq signal from biological replicates at each developmental stage was used. H3K9me3 signals represent Input-subtracted tracks, and transcription factor ChIP–seq signals represent IgG control–subtracted tracks. The same processed signal tracks used in Fig. 2B-2D were used for IGV visualization in Fig. 2E. All heatmaps are sorted based on H3K9me3 signal intensity in hepatoblasts. Same Y and Z (heatmap intensity) scales are used in Fig.2B and Fig.2C.

To determine whether H3K9me3 enrichment corresponds to functional repression, we analyzed nascent transcription using GRO-seq data from adult liver, as enhancer RNA expression serves as a reliable indicator of enhancer activity (Rahnamoun et al. 2020; Sartorelli and Lauberth 2020). Adult islet- and cortex-specific enhancers—regions that remain H3K9me3-marked in hepatocytes—exhibited minimal nascent RNA signal and low chromatin accessibility. In contrast, adult liver-specific enhancers displayed robust enhancer-associated transcription and open chromatin, features that anti-correlate with H3K9me3 enrichment (Fig. 2D). These findings indicate that persistent H3K9me3 marks correlate with transcriptional silencing of lineage-inappropriate enhancers.

Consistent with this model, hepatocyte lineage–determining transcription factors, including FOXA2, HNF4α, and members of the C/EBP family, preferentially occupied adult liver–specific enhancers, but not adult islet- and cortex-specific enhancers, that undergo H3K9me3 remodeling in differentiated liver (Fig. 2E, S2B). Together, these findings support a model in which liver-specific enhancer elements are maintained in a repressed chromatin state marked by H3K9me3 in undifferentiated progenitors and acquire transcriptional competence as H3K9me3 levels decline during differentiation.

### Lineage-specific enhancers are highly enriched for H3K9me3 in multiple tissue stem and progenitor populations

To assess whether enrichment of H3K9me3 at lineage-specific enhancers represents a general feature of tissue stem and progenitor cells, we analyzed H3K9me3 ChIP–seq datasets across mouse pancreatic β-cell differentiation, spanning stages from definitive endoderm to mature β cells (Fig. 3A). Adult islet-specific enhancers were defined based on previously published datasets (Tennant et al. 2013) (Table S7). Genomic annotation revealed that adult islet-specific enhancers are predominantly located in intergenic and intronic regions, similar to adult liver–specific enhancers (Fig. S3A). Consistent with our observations during hepatocyte differentiation, adult islet-specific enhancers were strongly enriched for H3K9me3 in pancreatic progenitor populations, indicating a repressed chromatin state (Fig. 3B). This enrichment was subsequently lost in immature and mature β cells, coinciding with PDX1 binding and increased chromatin accessibility (Fig. 3C; Fig. S3B).

**Fig. 3.**
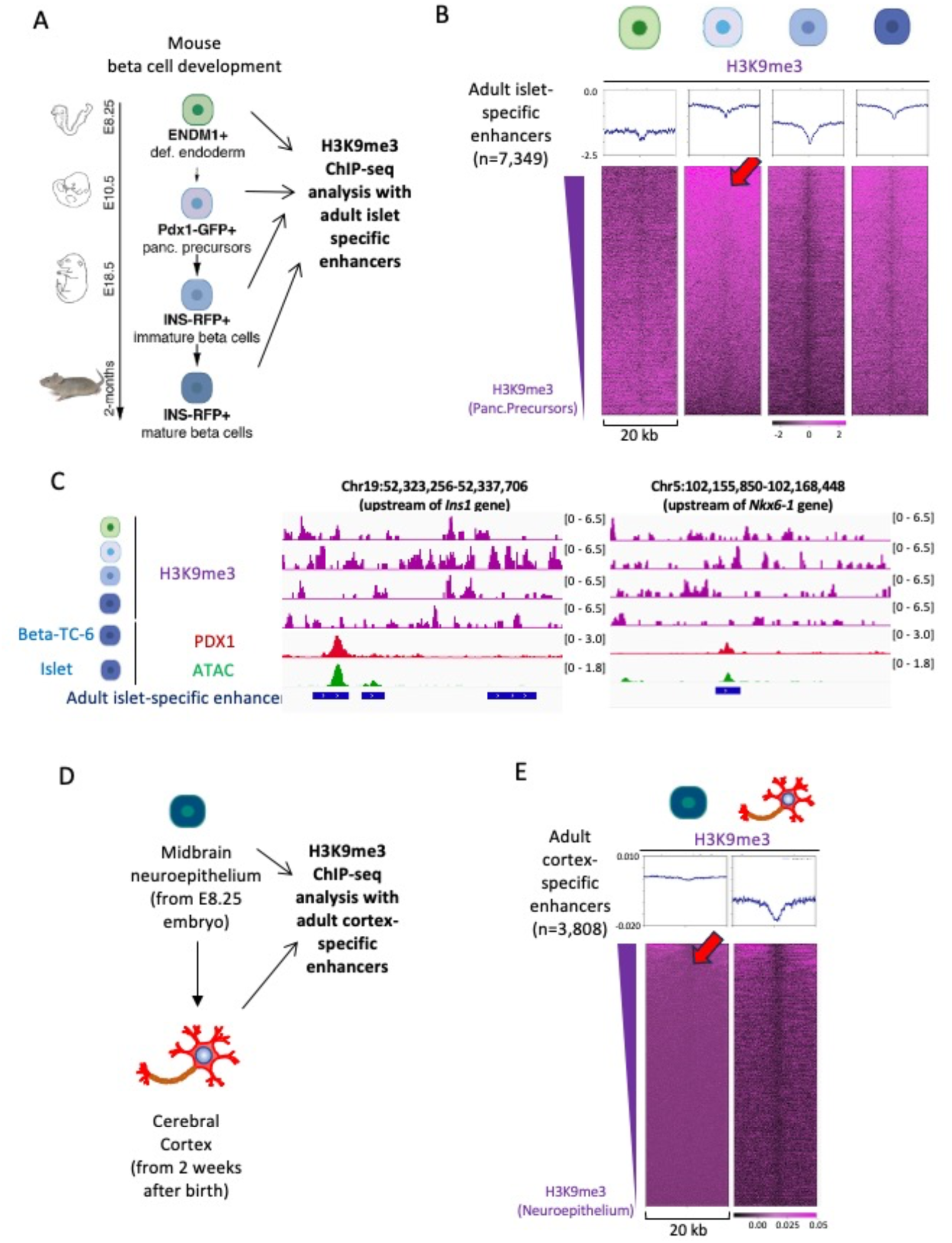
Lineage-specific enhancers in pancreas and cortex exhibit progenitor-enriched H3K9me3. A. Schematic illustration of the developmental stages and cell types analyzed in panels B and C. B. H3K9me3 heatmaps centered on adult islet-specific enhancers across pancreatic differentiation stages. C. Genome browser view of the β cell–specific *Ins1* and *Nkx6-1* loci showing H3K9me3 dynamics during pancreatic differentiation alongside PDX1 binding and chromatin accessibility in differentiated Beta TC-6 cells and islets. D. Schematic illustration of the developmental stages and cell types analyzed in panel E. E. H3K9me3 heatmaps centered on adult cortex-specific enhancers across cortical differentiation stages. For heatmap visualization, averaged H3K9me3 ChIP–seq signal from biological replicates at each developmental stage was used when available. For the adult cortex stage, a single available dataset was used. Heatmaps are sorted according to H3K9me3 signal intensity in the corresponding progenitor population.

To extend the analysis to a distinct developmental lineage, we examined H3K9me3 ChIP–seq datasets across mouse cortex development, from neuroepithelial progenitors to differentiated cortex (Fig. 3D). Genomic annotation revealed that adult cortex-specific enhancers are also predominantly located in intergenic and intronic regions (Fig. S3A). Consistent with the liver and pancreatic lineages, adult cortex-specific enhancers were strongly enriched for H3K9me3 in undifferentiated neuroepithelial cells, indicating a repressed chromatin state in progenitors. H3K9me3 levels were markedly reduced upon cortical differentiation (Fig. 3E; Fig. S3C).

Adult liver-specific enhancers were marked by H3K27me3 in definitive endoderm and mature β cells (Fig. S3D), similar to our observations for H3K9me3 (Fig. 2B, 2C). Likewise, adult islet-specific enhancers are marked by H3K27me3 in definitive endoderm and differentiated hepatocytes (Fig. S3D, S3E), similar to our observations for H3K9me3 (Fig. 2C, 3B). These results indicate that H3K9me3 and H3K27me3 appear cooperatively at tissue-specific enhancers in tissue stem and progenitor cells across multiple developmental lineages, as previously reported in mouse embryonic fibroblasts (Fukuda et al. 2023).

### Identification of cofactors associated with enhancer-like regulatory regions in the adult *liver*

To investigate factors associated with dynamic H3K9me3 remodeling at adult liver-specific enhancers, we focused on protein complexes recruited by the hepatocyte lineage–determining transcription factor HNF4α. Previous studies have shown that HNF4α co-recruits HDAC3 and PROX1 in differentiated liver, and mass spectrometry analysis of HDAC3-associated proteins identified multiple cofactors, including NCOR1 and NR3C1 (Armour et al. 2017). Analysis of ChIP-seq datasets for these cofactors revealed preferential binding to adult liver-specific enhancers in differentiated liver (Fig. 4A; Fig. S4A).

**Fig. 4.**
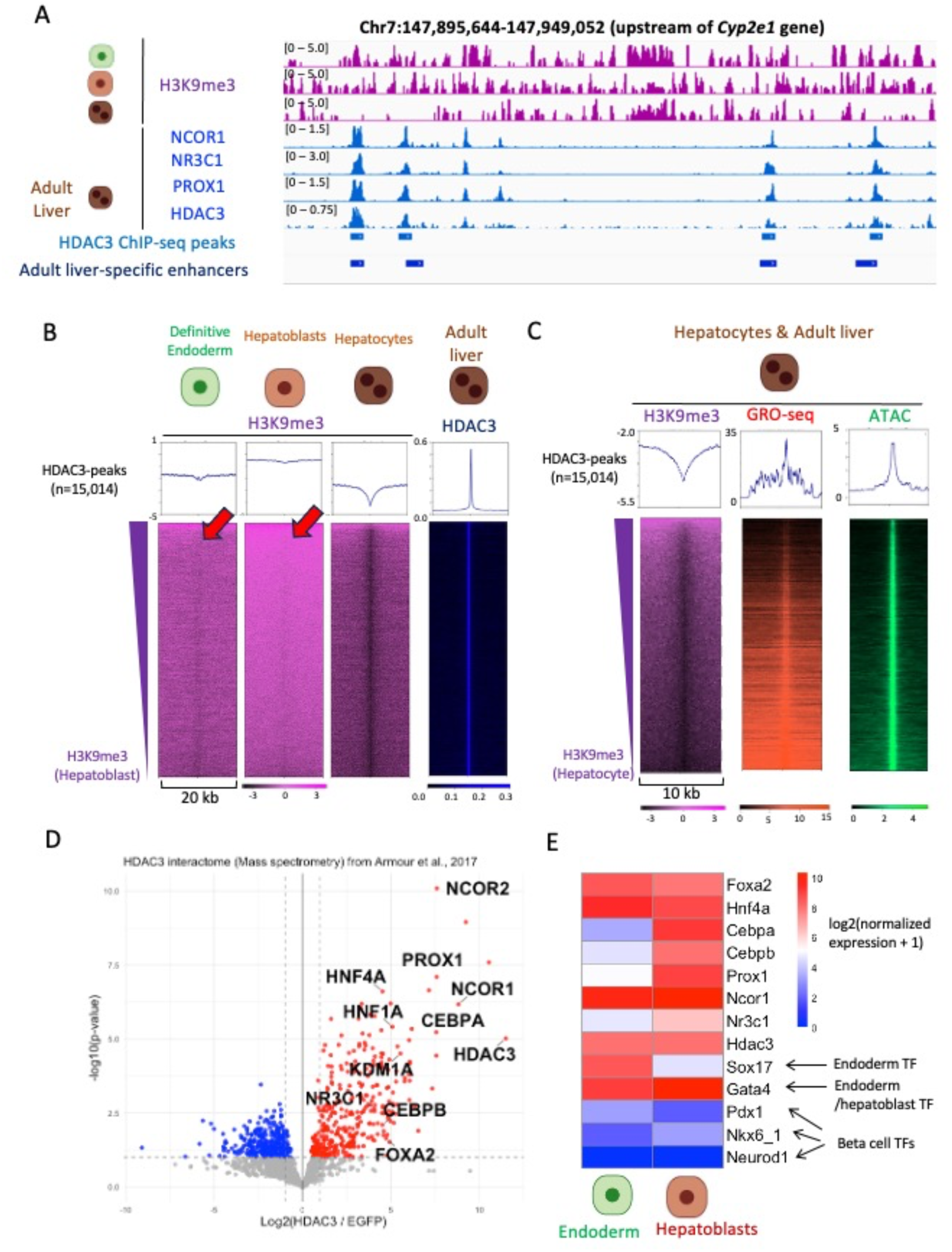
Identification of factors associated with enhancer-like regulatory regions in the adult liver. A. Genome browser view of the upstream region of the *Cyp2e1* locus showing HDAC3 and associated cofactors binding to enhancer-like regulatory regions in differentiated liver that exhibit dynamic H3K9me3 remodeling during liver development. B. H3K9me3 and HDAC3 heatmaps centered on HDAC3 binding sites across liver developmental stages. C. H3K9me3, GRO-seq, and ATAC-seq heatmaps in adult liver centered on HDAC3 peaks. Positive- and negative-strand GRO-seq signals were merged prior to analysis. D. Reanalysis of the HDAC3 interactome (mass spectrometry data from Armour et al., 2017) identifying hepatocyte lineage–determining transcription factors associated with HDAC3. E. Heatmap showing expression of hepatocyte lineage–determining transcription factors and cofactors in definitive endoderm and hepatoblasts based on published RNA-seq data (Nicetto et al., 2019). For heatmap visualization, averaged H3K9me3 ChIP–seq signal from biological replicates at each developmental stage was used. H3K9me3 signals represent Input-subtracted tracks, and transcription factor ChIP–seq signals represent IgG control–subtracted tracks. The same processed signal tracks were used for IGV visualization and quantitative analyses in Fig. 4A- 4C. The heatmap in Fig. 4B is sorted based on H3K9me3 signal intensity in hepatoblasts, and the heatmap in Fig. 4C is sorted based on H3K9me3 signal intensity in hepatocytes. Same Y and Z (heatmap intensity) scales as in Fig.2 are used in Fig. 4B.

To further characterize the chromatin features of HDAC3-bound regions, we examined H3K9me3 ChIP-seq profiles centered on HDAC3 binding sites. Notably, HDAC3-associated regions exhibited adult liver-specific enhancer-like H3K9me3 dynamics, showing strong enrichment of H3K9me3 in undifferentiated definitive endoderm and hepatoblasts, followed by marked loss of H3K9me3 in differentiated hepatocytes (Fig. 4B). Consistent with this enhancer-like behavior, GRO-seq analysis demonstrated that HDAC3 peaks were associated with robust enhancer-associated transcription and open chromatin, features that anti-correlate with H3K9me3 enrichment, similar to adult liver-specific enhancers (Fig. 4C). Thus, HDAC3-bound regions appear to be activated like liver-specific enhancers following H3K9me3 removal.

Given that HDAC3 preferentially associates with adult liver-specific enhancers, we reasoned that HDAC3-associated mass spectrometry data could be leveraged to identify factors functionally linked to adult liver-specific enhancers. Consistent with this idea, several transcription factors previously shown to be enriched at adult liver-specific enhancers (Fig. 2E, S2B), including FOXA2, HNF4α, and members of the C/EBP family, were co-precipitated with HDAC3 (Fig. 4D). In agreement with this finding, ChIP–seq peaks of these transcription factors were frequently detected in proximity to HDAC3 binding sites (Fig. S4B). As expected, the enhancer-associated lysine-specific histone demethylase KDM1A (LSD1) was also detected in HDAC3-containing complexes (Fig. 4D). In contrast, H3K9me3 demethylases of the KDM4 family were not detected in HDAC3 immunoprecipitates, indicating that KDM4 proteins are unlikely to be directly involved in H3K9me3 removal at adult liver-specific enhancers during hepatocyte differentiation.

FOXA1/2, HNF4α, and members of the C/EBP family are known to be expressed prior to or early during hepatic lineage commitment (Ang et al. 1993; Duncan et al. 1994; Lee et al. 2005; Westmacott et al. 2006). To examine the temporal expression of transcription factors identified in the HDAC3 interactome, we analyzed previously published RNA-seq datasets from undifferentiated definitive endoderm and hepatoblasts (Nicetto et al. 2019). Notably, multiple transcription factors, including *Foxa2* and *Hnf4α*, as well as cofactors such as *Ncor1* and *Hdac3*, were already expressed at the definitive endoderm stage at levels comparable to the endodermal marker *Sox17*. In addition, other transcription factors and cofactors, including members of the *C/EBP* family, *Prox1*, and *Nr3c1*, were expressed in hepatoblasts at levels comparable to the hepatoblast marker *Gata4* (Fig. 4E).

Together, these observations suggest that H3K9me3 enrichment at adult liver–specific enhancers in undifferentiated progenitor populations restrains premature enhancer activation, even though hepatocyte lineage–determining transcription factors are already expressed. This raises the possibility that H3K9me3 may function to limit transcription factor access to lineage-specific enhancers prior to differentiation.

### H3K9me3 differentially restricts pioneer factor access to distinct classes of adult liver-specific enhancers in hepatoblasts

Previous studies have shown that the pioneer factor FOXA2 binds to adult liver-specific enhancers and enables subsequent liver-determining transcription factor binding to the enhancers (Iwafuchi-Doi et al. 2016). To study the importance of H3K9me3 for liver-determining transcription factor binding during liver development, we tested the behavior of H3K9me3 around FOXA2 and HNF4α binding loci during liver development. To do this, we compared FOXA2 and HNF4α ChIP-seq data and ATAC-seq data between hepatoblasts and hepatocytes. Since the datasets were generated by different laboratories in independent studies, ChIP/ATAC-seq data were normalized using peaks at the *Albumin* enhancer, where these transcription factors are known to bind in both hepatoblasts and hepatocytes (Cirillo et al. 2002) (Fig. S5A). We used these normalized ChIP/ATAC-seq data for subsequent analyses.

We first categorized adult liver-specific enhancers based on FOXA2 binding peaks observed around adult liver-specific enhancers (5 kb window size) (Fig. 5A, top). As expected, the percentage of enhancers in which hepatoblast-specific FOXA2 peaks are observed is very small (1.3% of total adult liver-specific enhancers; categorized as class i). The percentage of enhancers in which FOXA2 peaks are commonly observed in both hepatoblasts and hepatocytes is 12% (categorized as class ii). The percentage of enhancers in which hepatocyte-specific FOXA2 peaks are observed is 39% (categorized as class iii). The total percentage of adult liver-specific enhancers categorized into class ii and iii is more than 50%, supporting that the pioneer factor FOXA2 is a key transcription factor for liver development, as reported in previous studies (Lee et al. 2005; Iwafuchi-Doi et al. 2016).

**Fig. 5.**
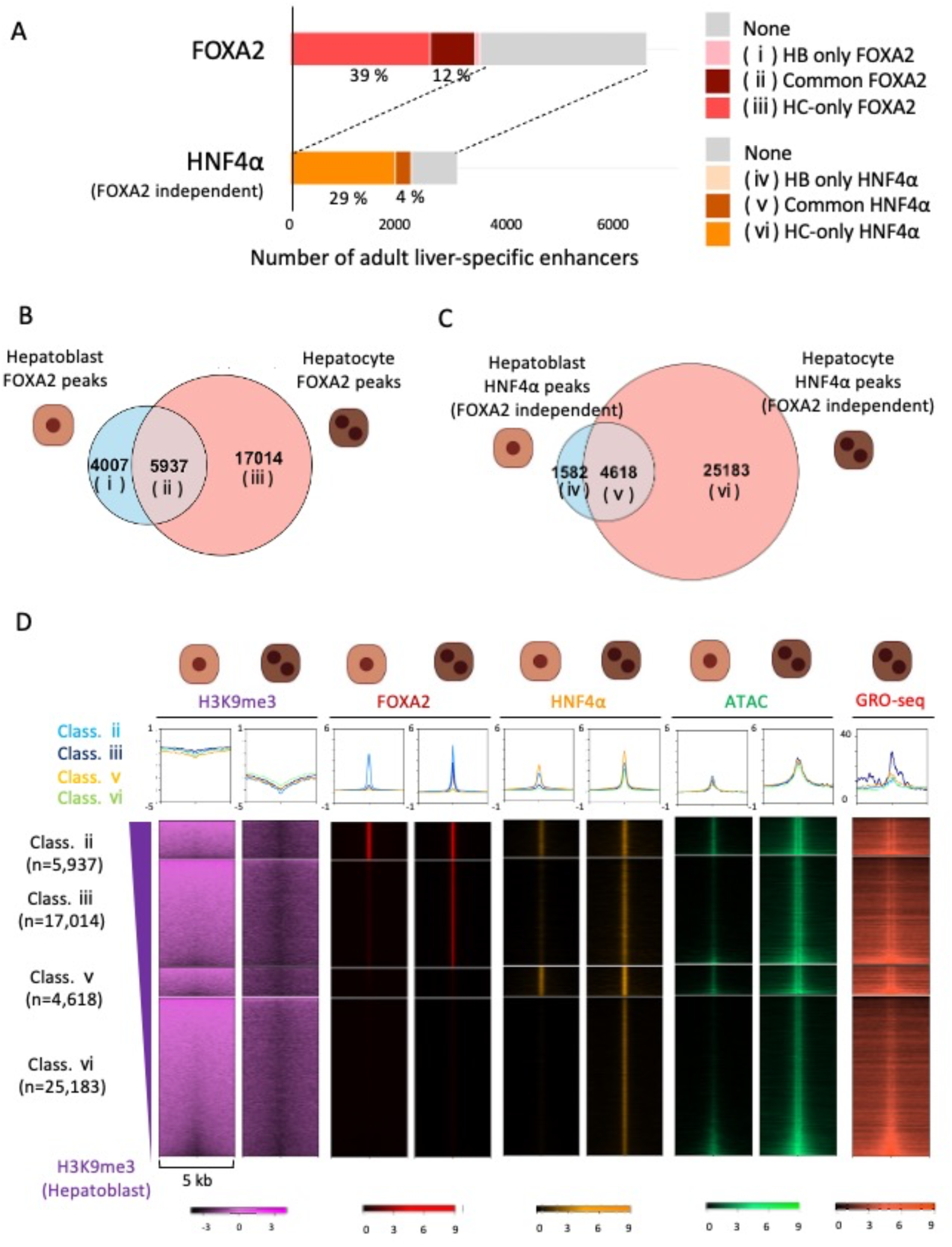
Lineage-determining transcription factor binding to adult liver–specific enhancers is constrained in hepatoblasts when these regions are marked by H3K9me3. A. Bar graph showing the percentage of adult liver-specific enhancers categorized based on FOXA2 and FOXA2-independent HNF4α peaks in hepatoblasts and hepatocytes. B. Venn diagram representing the total number of FOXA2 peaks in hepatoblasts and hepatocytes. C. Venn diagram representing the total number of FOXA2-independent HNF4α peaks in hepatoblasts and hepatocytes. D. H3K9me3, FOXA2, HNF4α, and chromatin accessibility (ATAC-seq) heatmaps in hepatoblasts and adult liver, and GRO-seq in adult liver, centered on class ii, iii, v, and vi. H3K9me3 ChIP-seq data were obtained from Liv2⁺ hepatoblasts at E10.5, and FOXA2 and HNF4α ChIP-seq data and ATAC-seq data were obtained from DLK1⁺ hepatoblasts at E14.5. For heatmap visualization, averaged H3K9me3 ChIP-seq signal from biological replicates at each developmental stage was used. H3K9me3 signals represent Input-subtracted tracks, and transcription factor ChIP-seq signals represent IgG control-subtracted tracks. Heatmaps are sorted based on H3K9me3 signal intensity in hepatoblasts.

Regarding adult liver-specific enhancers in which FOXA2 peaks were not observed, we reasoned that they are regulated by another key liver transcription factor, HNF4α, independently of FOXA2 (5 kb window size) (Fig. 5A, bottom). Among these FOXA2-negative adult liver-specific enhancers, the percentage in which hepatoblast-specific HNF4α peaks are observed is very small (less than 1.0% of total adult liver-specific enhancers; categorized as class iv). The percentage in which HNF4α peaks are commonly observed in both hepatoblasts and hepatocytes is 4% of total adult liver-specific enhancers (categorized as class v). The percentage in which hepatocyte-specific HNF4α peaks are observed is 29% (categorized as class vi). These analyses indicate that class iii and vi represent the major classes, whereas class ii and v represent minor classes of adult liver-specific enhancers.

To extend the analysis, total FOXA2 peaks were also categorized into class i to iii, similar to Fig. 5A. 4,007 and 17,014 FOXA2 peaks were observed as hepatoblast-specific and hepatocyte-specific FOXA2 peaks, respectively (Fig. 5B). The number of FOXA2 peaks commonly observed in both hepatoblasts and hepatocytes is 5,937 and they were categorized into class i to iii. Similarly, total FOXA2-independent HNF4α peaks were categorized into classes iv to vi, similar to Fig. 5A. 1,582 and 25,183 FOXA2-independent HNF4α peaks were observed as hepatoblast-specific and hepatocyte-specific FOXA2-independent HNF4α peaks, respectively (Fig. 5C). The number of FOXA2-independent HNF4α peaks commonly observed in both hepatoblasts and hepatocytes is 4,618 and they were categorized into classes iv to vi (All the lists are provided in Table S10 and S11).

To examine H3K9me3 behavior around classes ii, iii, v, and vi, in which adult liver-specific enhancers are included, we generated heatmaps. Dynamic loss of H3K9me3 was observed around these classes with concomitant changes in chromatin accessibility (Fig. 5D), similar to the heatmaps generated for all adult liver-specific enhancers (Fig. 2B). FOXA2 and HNF4α binding to classes iii and vi is restricted by H3K9me3 in hepatoblasts. Notably, FOXA2 and HNF4α also bind to a substantial number of H3K9me3-marked enhancers in hepatoblasts—encompassing approximately 5,937 FOXA2 peaks and 4,618 HNF4α peaks in classes ii and v, respectively—suggesting that pioneer factor engagement with H3K9me3-marked chromatin is not absolutely prohibited and may reflect a state of developmental competence at these sites. The average chromatin accessibility levels of these classes in hepatoblasts did not differ, suggesting that different co-factors binding to each category may account for the difference. Adult liver-specific enhancers are observed in these classes with H3K9me3 and chromatin accessibility dynamics during liver development (Fig. S5B). As controls, we also generated heatmaps focusing on classes i and iv, in which adult liver-specific enhancers are not observed. Dynamic changes in H3K9me3 and chromatin accessibility were not observed in these categories, despite FOXA2 and HNF4α binding being observed in hepatoblasts (Fig. S5C). Transcriptional activity around these control classes in hepatocytes was lower than that of both the major and minor classes (Fig. 5D, Fig. S5C). Taken together, the results indicate that pioneer factor binding and chromatin opening during liver development are modulated not only by H3K9me3 state, but can also by modulated by co-factors that bind to each category during liver development.

## Discussion

We have described chromatin-based features of tissue stem and progenitor cells in which lineage-specific promoters and enhancers are preferentially marked by H3K9me3 prior to differentiation. By integrating enhancer annotation with HDAC3-associated mass spectrometry datasets, we identified transcription factors and cofactors critical for hepatocyte specification as components associated with enhancer-linked regulatory regions connected to adult liver-specific enhancers. Notably, many of these factors are already expressed in undifferentiated definitive endoderm and hepatoblasts. By integrating ChIP-seq datasets across different developmental stages, adult liver-specific enhancers were categorized into distinct classes and H3K9me3 was shown to contribute to maintaining the undifferentiated state in tissue stem and progenitor populations by distinct mechanisms. In the major classes, H3K9me3 restricts premature transcription factor engagement and prevents inappropriate enhancer activation by establishing a compact chromatin state. In the minor classes, which together encompass thousands of binding sites, transcription factors engage H3K9me3-marked enhancers in hepatoblasts, yet the chromatin remains compact, suggesting that H3K9me3 contributes to maintaining the undifferentiated state through mechanisms that act downstream of or in parallel with transcription factor binding (Fig. 6).

**Fig. 6.**
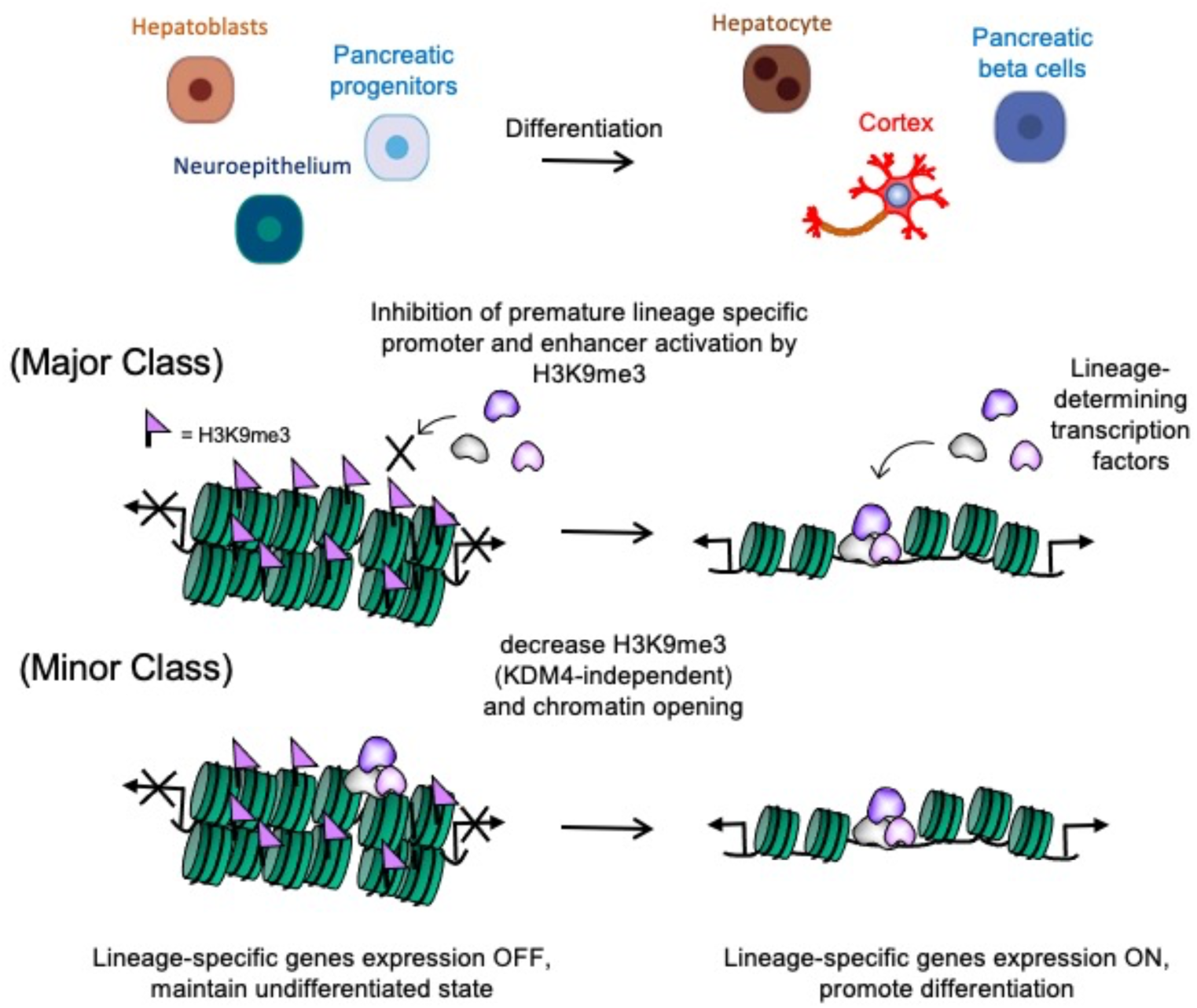
Conceptual model illustrating KDM4-independent modulation of H3K9me3 at lineage-specific enhancers during differentiation from tissue stem and progenitor cells. H3K9me3 restrains lineage-specific enhancers in progenitor cells by limiting premature engagement of lineage-determining transcription factors, independently of KDM4-mediated demethylation, until differentiation cues trigger chromatin remodeling and enhancer activation.

Given that lineage-specific enhancers are marked by H3K9me3 across multiple tissue stem and progenitor populations, extrinsic signaling pathways and their downstream effectors may contribute to the maintenance of the undifferentiated state by regulating H3K9me3 enrichment at these enhancers. Although H3K9me3 demethylation is often attributed to members of the KDM4 family (Das et al. 2014; Sankar et al. 2020), proteomic and chromatin analyses suggest that KDM4 proteins are unlikely to be directly involved in the dynamic regulation of H3K9me3 at adult liver-specific enhancers during hepatocyte differentiation. This observation raises the possibility that H3K9me3 dynamics at lineage-specific enhancers are governed by mechanisms other than active demethylation by the KDM4 family, such as replication-coupled dilution or histone acetyltransferase recruitment by transcription factors.

In the final part of this study, we showed that the binding of FOXA2 and HNF4α to the major enhancer classes (class iii and vi in Fig. 5D) is restricted in hepatoblasts when these regions remain enriched for H3K9me3. This finding is consistent with previous studies reporting that ectopic expression of many but not all pioneer factors fails to overcome H3K9me3-marked chromatin barriers in fibroblasts (Soufi et al. 2012; Donaghey et al. 2018; Katznelson et al. 2026). Our data extend this concept to developmental contexts, suggesting that H3K9me3 can similarly constrains pioneer factor engagement at lineage-specific enhancers in tissue stem and progenitor cells. At the same time, we also observed the binding of FOXA2 and HNF4α to the minor repressed classes (class ii and v in Fig. 5D) in hepatoblasts. This finding supports a recent study which showed that FOXA2, but not HNF4α, can engage a small subset of H3K9me3-marked heterochromatic loci when ectopically expressed (Katznelson et al. 2026), suggesting that pioneer factor binding to H3K9me3-enriched regions is not absolutely prohibited but may depend on dosage and chromatin context. Taken together, these findings raise the possibility that pioneer factor access to H3K9me3-marked enhancers is modulated by additional cofactors under physiological developmental conditions. For example, GATA4—although not detected in the HDAC3 mass spectrometry dataset—is known to bind the *Albumin* enhancer in hepatoblasts (Bossard and Zaret 1998; Bossard and Zaret 2000) and may contribute to this tissue-specific enhancer classification. This is further supported by a previous study showing that chromatin opening in vivo by FOXA2 can require collaboration with additional transcription factors (Cernilogar et al. 2019). In contrast to the major and minor classes, pioneer factor FOXA2 binding was observed in class i, where dynamic changes in H3K9me3 and chromatin accessibility were not observed (Fig. S5C). This finding supports recent studies showing that pioneer factors contribute to the loss of cellular identity by recruiting repressor complexes (Chronis et al. 2017; Katsuda et al. 2024; Matsui et al. 2024). The distinct outcomes of FOXA2 binding across enhancer classes may therefore reflect differences in the co-factors present at each class, ranging from repressor complexes in class i to activating factors in the major and minor classes.

Collectively, these observations suggest that H3K9me3 does not simply act as a static heterochromatic barrier but rather functions as a dynamic and context-dependent regulatory mechanism that modulates pioneer factor access during lineage specification. This dynamic regulation may provide a molecular mechanism to fine-tune developmental timing while preserving lineage fidelity.

## Experimental procedures

### Identification of adult liver-enriched genes and their promoters

Genes belonging to Cluster #13, characterized by exclusive expression in the differentiated liver during development based on published bulk RNA-seq data (Nicetto et al. 2019), were defined as adult liver-enriched genes. To define their promoter regions, genomic coordinates for the mouse genome (mm9/NCBI37) were retrieved using the *TxDb.Mmusculus.UCSC.mm9.knownGene* and *org.Mm.eg.db* R/Bioconductor packages to ensure consistency with previously published datasets. Promoters were defined as the regions extending 500 bp upstream and 500 bp downstream of the transcription start site (TSS). Genes were filtered to include only those with valid Entrez ID mappings and annotated coordinates within the mm9 assembly; genes with updated symbols or those lacking genomic records in the mm9 database were excluded from the final BED files. The resulting coordinates were exported in BED format for subsequent analyses. The list of adult liver-enriched genes and their promoters used in this study are provided in Supplemental Tables S1 and S2, respectively.

### Identification of adult liver-specific genes and their promoters

Adult liver-specific promoters were defined as the subset of liver-enriched gene promoters overlapping by at least 1 bp with adult liver-specific RNA Pol II peaks (3 kb window size; Iwafuchi-Doi et al., 2016). These promoters and their associated genes were identified using the R/Bioconductor packages *GenomicRanges* and *rtracklayer*. Genomic coordinates were mapped to the mouse reference genome (mm9; NCBI37) using the *TxDb.Mmusculus.UCSC.mm9.knownGene* annotation package to ensure consistency with previously published datasets. Entrez IDs were converted to official Gene Symbols via *org.Mm.eg.db*. The mapping of 147 identified promoter regions resulted in 143 unique genes, accounting for cases where multiple promoter peaks were assigned to a single gene.

The list of adult liver-specific genes and their promoters used in this study are provided in Supplemental Tables S3 and S4, respectively.

### Identification of β-cell and neuron-specific genes and their promoters

Previously reported gene lists for differentiated β-cell-specific (Benner et al. 2014) and neuron-specific genes (Sanosaka et al. 2017) were utilized. To identify the promoter regions of these genes, genomic coordinates for the mouse genome (mm9/NCBI37) were retrieved using the *TxDb.Mmusculus.UCSC.mm9.knownGene* and *org.Mm.eg.db* R packages to ensure consistency with previously published datasets. Promoters were defined as the regions extending 500 bp upstream and 500 bp downstream of the TSS. Genes were filtered to include only those with valid Entrez ID mappings and annotated coordinates within the mm9 assembly; genes with updated symbols or those lacking genomic records in the mm9 database were excluded from the final BED files. The resulting coordinates were exported in BED format for subsequent analyses. The list of β-cell and neuron-specific genes and their promoters used in this study are provided in Supplemental Tables S5 and S6, respectively.

### Generation of lineage-specific enhancer lists

Adult liver-specific enhancer peaks (1kb window size; GSE29184) that overlap by at least 1 bp with DNase I hypersensitive sites peaks (1kb window size; GSM1014195) were defined as adult liver-specific enhancers in this study.

Adult islet-specific enhancers were obtained from a previously published dataset (Tennant et al. 2013). Adult cortex-specific enhancer peaks (1kb window size; GSE29184) that overlap by at least 1 bp with ATAC-seq positive peaks (1kb window size; GSM4905596) were defined as adult cortex-specific enhancers. To generate adult liver-and cortex-specific enhancer lists, we used previously published enhancer datasets identified by Shen and colleagues based on ChIP–seq data for p300, H3K4me1, and H3K4me3 in adult mouse liver and cortex (ENCODE accession GSE29184) (Shen et al. 2012).

Genomic annotation of enhancer sets was performed using the R package *ChIPseeker*, and genomic distribution plots were generated using *seqplot* to assess the proportion of peaks located in intergenic, intronic, promoter, and exonic regions. Genomic annotation was performed using the mm9 TxDb annotation database (TxDb.Mmusculus.UCSC.mm9.knownGene). All analyses were performed using the mm9 mouse genome assembly to maintain consistency with previously published datasets. All enhancer lists used in this study are provided in Supplemental Table S7.

### Alignment of H3K9me3/H3K27me3 ChIP-seq data

For H3K9me3/H3K27me3 ChIP–seq analyses, biological replicates obtained from previous studies (Nicetto et al. 2019) were merged at the BAM level when available to increase signal robustness. Input-subtracted signal tracks were generated, and the resulting spike-in-normalized BigWig (mm9) files (Nicetto et al. 2019) were used for all quantitative and visual analyses. When only a single dataset was available, that dataset was processed using the same pipeline. Processed BigWig files were visualized using IGV for representative loci and used for comparative analyses across developmental stages. Corresponding GEO accession numbers are listed in Supplemental Table S8. All analyses were performed using the mm9 mouse genome assembly to maintain consistency with previously published datasets.

### RNA-seq heatmap analysis for lineage-specific genes

To visualize the expression patterns of the identified gene sets, heatmap analysis was performed using the pheatmap package in R. Normalized expression values for the selected genes were retrieved from Supplemental Table S13 of (Nicetto et al. 2019). Before visualization, lineage-specific gene lists were integrated, and overlapping genes were assigned to a single category following a priority hierarchy (adult liver-specific > - cell enriched > neuron-specific) to ensure row uniqueness. Genes that were absent from the reference dataset or exhibited zero variance across the selected lineages were excluded. This filtering process resulted in final sets of 143 adult liver-specific, 90 β-cell-specific, and 454 neuron-specific genes (all listed in Supplemental Table S9). For effective comparison of expression dynamics, the data were subjected to gene-wise Z-score transformation. Columns were manually ordered to reflect the hepatic developmental trajectory: definitive endoderm, hepatoblasts, and adult hepatocytes. The heatmap was presented without row clustering to maintain the predefined hierarchical group order.

### Gene Ontology enrichment analysis

Gene Ontology (GO) enrichment analysis was performed on the identified adult liver-specific genes using the R package clusterProfiler. Gene symbols were mapped to Entrez IDs using the org.Mm.eg.db database. Functional enrichment was assessed for the Biological Process (BP) category. Statistical significance was determined using a one-sided hypergeometric test, and p-values were adjusted for multiple testing using the Benjamini-Hochberg (BH) method. A significance threshold of adjusted p-value < 0.05 (or q-value < 0.05) was applied for the identification of enriched GO terms.

### Transcription factor/cofactor ChIP-seq data and ATAC-seq data

BigWig files of aligned transcription factor and cofactor ChIP–seq and ATAC-seq datasets were downloaded from ChIP-Atlas (Oki et al. 2018; Zou et al. 2022; Zou et al. 2024). For all transcription factor ChIP–seq datasets, IgG control–subtracted signal tracks were generated using deepTools (bigwigCompare, --operation subtract) and used for visualization and quantitative analyses. Peak files for transcription factor ChIP–seq were obtained from ChIP-Atlas (q < 1E-05) and used for analyses. Corresponding GEO accession numbers are listed in Table S8. All analyses were performed using the mm9 mouse genome assembly to maintain consistency with previously published datasets.

### GRO-seq heatmap analysis

RPM-normalized BigWig files of aligned GRO-seq data from adult liver were downloaded from GEO (accession numbers listed in Table S8). For each biological replicate, positive- and negative-strand signals were merged to generate total nascent transcription signal. The merged tracks from two biological replicates were then averaged and used for all heatmap and genome browser visualizations. All analyses were performed using the mm9 mouse genome assembly to maintain consistency with previously published datasets.

### ChIP-seq heatmap analysis

Heatmaps were generated using deepTools (Ramirez et al. 2016). BigWig files were used to compute signal enrichment over defined genomic regions using computeMatrix, and heatmaps were visualized with plotHeatmap.

### Mass spectrometry data

HDAC3-associated mass spectrometry data (Armour et al. 2017) were analyzed using R (version 4.2.2). Protein-level quantitative values were imported into R and log2 fold changes between HDAC3 immunoprecipitation and control samples were calculated. Statistical significance was assessed using the p-values provided in the mass spectrometry output. Volcano plots were generated using the ggplot2 package, with log2 fold change plotted against −log10 (p-value). Selected proteins of interest were annotated using ggrepel.

### Normalization and quantitative scaling of FOXA2 and HNF4α ChIP/ATAC-seq data

To ensure quantitative comparability of FOXA2, HNF4α occupancy and chromatin accessibility across liver developmental stages, BigWig tracks for FOXA2, HNF4α, and ATAC-seq were obtained from ChIP-Atlas (Oki et al. 2018; Zou et al. 2022; Zou et al. 2024), where signal intensities were pre-normalized to the number of reads per million mapped reads (RPM). To further refine the quantitative comparison between hepatoblast (HB) and adult hepatocyte (HC) stages, we applied a localized scaling method centered on the *Alb* (Albumin) and *Afp* (Alpha-fetoprotein) loci. These loci served as internal biological controls: the *Alb* locus, which maintains high accessibility and robust factor binding in both stages (Cirillo et al. 2002), was used to calibrate the upper dynamic range of the signals. Conversely, the *Afp* locus, which exhibits high enrichment in HB but marked reduction in HC, was used to verify the sensitivity of the scaling to stage-specific transitions. Practically, BigWig files were converted to bedGraph format using bigWigToBedGraph, and signal intensities were linearly transformed by multiplying the scores by calculated scaling factors. The resulting scores were then re-converted to the BigWig format for subsequent analyses. These factors were adjusted such that the peak heights at these conserved and stage-specific control regions were consistent with their expected biological activity, allowing for the direct comparison of signal enrichment at other liver-specific enhancers.

### FOXA2 and HNF4α binding dynamics analysis

To characterize the global binding shifts of FOXA2 and HNF4α during liver development, peak files for FOXA2 and HNF4α ChIP–seq from embryonic hepatoblast (HB) and adult hepatocyte (HC) were obtained from ChIP-Atlas (q < 1E-05) (Oki et al. 2018; Zou et al. 2022; Zou et al. 2024). FOXA2 peaks were categorized into HB-only (class i), Common (class ii), HC-only (class iii) groups using a 3 kb window. For HNF4α, we specifically defined “FOXA2-independent peaks” by excluding any signals overlapping with FOXA2 occupancy within a 3 kb window in the respective stage. These FOXA2-independent HNF4α peaks were further categorized into HB-only (class iv), Common (class v), HC-only (class vi) groups using strict mutual-exclusion criteria, where a peak was considered stage-specific only if no HNF4α occupancy was detected within a 3 kb window in the opposing developmental stage. Proportional Venn diagrams were generated using the eulerr package. The final classified peak lists are provided in Supplemental Table S10.

### Classification of adult liver-specific enhancers

Adult liver-specific enhancers were categorized into six functional classes (class i–vi) based on the occupancy of the FOXA2 and FOXA2-independent HNF4α peak groups defined above. An enhancer was considered “occupied” if a peak was present within a ±2.5 kb window from the enhancer center. Enhancers were first grouped by FOXA2 occupancy: HB-only (class i), Common (class ii), and HC-only (class iii). The remaining FOXA2-independent enhancers were then sub-classified based on their overlap with the previously defined FOXA2-independent HNF4α peak groups: HB-only (class iv), Common (class v), and HC-only (class vi). All genomic interval manipulations were performed using GenomicRanges and rtracklayer, and the final classified adult liver-specific enhancer lists are provided in Supplemental Table S11.

## Acknowledgments

We thank current and former Zaret lab members for technical advice and discussion. And we also thank Dr. Oki for developing ChIP-Atlas.

## Author contributions

Conceptualization: K.I., K.Z. Investigation: K.I., G.D. Formal analysis: K.I., G.D.

Writing - Original Draft: K.I., K.Z. Writing – Review & Editing: All authors

Supervision and Funding acquisition: K.I., K.Z.

